# Categorizing Ideas about Systematics: Alternative Trees of Trees, and Related Representations

**DOI:** 10.1101/079483

**Authors:** János Podani, David A. Morrison

## Abstract

This study is an attempt to expand a previous survey by Fisler and Lecointre (FL) for systematizing ideas on the use of the tree metaphor in classification, as expressed by various historically important figures in their writings. FL used a cladistic approach to analyze their data, as employed in biological classification. We supplement this analysis here using several methods of multivariate data exploration, producing a UPGMA dendrogram, a minimum spanning tree, a neighbor joining additive tree, a plexus graph, a phylogenetic network, and two multidimensional scaling ordinations of the same data used by FL. We confirm the validity of many of FL’s smaller clusters of writings, and revealed a new 3-group categorization undetected by the previous study. These three groups largely correspond to *Classifiers*, who did not consider evolution for historical reasons or on purpose, *Non-analytical evolutionists*, who recognized evolution but with a more or less naïve attitude towards the temporal change of life, and *Modelers*, with more explicit views on evolutionary processes, often applying objective mathematical tools for exploring the past and present of organismal diversity. Some scientists were difficult to assign to any group unambiguously, including J.W. von Goethe, who takes a unique position in the history of biology, and, to a lesser extent, E. Mayr and G.G. Simpson, the leaders of the gradist school of systematics. We argue that cladistic methods are insufficient by themselves, notably in situations where there are no obvious ancestor-descendant relationships underlying the development of the objects being analyzed.

## 1 Introduction

In historical accounts of science, it is a common practice to assign labels to members of various intellectual schools, such as Pythagoreans, Essentialists, Darwinists, or Popperians. These categories are determined in most cases based on a fairly subjective basis, without formal quantitative analyses of the views expressed by designated representatives of these schools. A noted exception is a recent paper by Fisler and Lecointre (2013; abbreviated hereafter as FL) who recognized that ideas in biology may be described in terms of many measurable variables simultaneously, allowing the possibility of objective comparisons.

FL selected 41 published works with the purpose to categorize ideas about “phylogenetic” trees and tree-based classifications. These writings encompass several centuries of scientific advancement, from A. Zaluziansky who worked late in the sixteenth century to P. Tassy whose ideas were summarized at the end of the twentieth century. Both written text and drawings were evaluated for 91 different variables that conceptualize the ideas being expressed. The resulting 41 × 91 data matrix was analyzed by the cladistic method of maximum parsimony (rooted in the works of Hennig 1966), in order to reveal “clades”^1^ of authors and to see how the different ideas were shared by their proponents in these groups. All conclusions were based on a hierarchical arrangement, a so-called “tree of trees”, generated as a single, unweighted consensus cladogram of 279 equally parsimonious results. The authors were able to identify well-known schools of systematics, and they also specified some groups that are apparently new to the history of biology.

We find that whereas their pioneering approach is extremely interesting and thought-provoking, it is remarkably one-sided. The cladistic analysis is used, in fact, for classificatory purposes, rather than for other possible relationships among the objects, such as direct ancestry or inter-object dissimilarity evaluated without imposing cluster structure on the data. However, many other methods are widely available in the statistical literature, for both classification and other forms of relationship. The cladistic method adopted belongs to one particular school of systematics, although other schools have also employed objective procedures for tree-making. Cladistic methods have been used to reveal relationships for objects with an evolutionary history, such as organisms (Hennig 1966; and his followers), languages (Rexová et al. 2003), archaeological specimens (O’Brien et al. 2001), music (Le Bomin et al. 2015) or even biblical scripts (Howe and Windram 2011). The fact that cladistics does not well fit the complex development of biological thought is admitted by Fisler and Lecointre themselves, who said that “the flow of ideas through times doesn’t behave like in biological entities”. Similarities between ideas are obviously not due to simple “inheritance” or “ancestry” and therefore cladograms may not be the best approach, and are definitely not the only appropriate representations of quantifiable structure in the data.

Phenetics, another school of systematics covered by Fisler and Lecointre’s study, however, uses a much wider range of procedures for evaluating similarities, revealing categories as well as visualizing results in various graphical forms, such as networks, dendrograms and ordinations. There are also network-generating procedures, which do have applications in phylogenetics and elsewhere (Morrison 2014), whose capabilities were explicitly ignored by FL. Other tree-generating methods and ordination procedures are effective summaries of multivariate data, but as such they will differ from each other depending on which aspects of the data are emphasized in the summary. These methods are therefore usually complementary, in that when they are considered together they can reveal patterns that are not necessarily obvious in any one data summary. So, it is best to use a combination of clustering, network and ordination methods in order to thoroughly explore any given multivariate data set.

Our approach here is explicitly one of exploratory data analysis (Tukey 1977). This methodology eschews the idea of testing formal hypotheses that can be stated a priori, but instead explores the data in a model-independent manner. Graphical representations of the data are an important part of data exploration (Ellison 2001), rather than formal statistical analyses. Exploratory data analysis is useful in any field of science, from anthropology through psychology to zoology, including phylogenetics (Morrison 2010), in which many objects are described in terms of many features or variables.

It is important to note that time is not explicitly incorporated into any of the multivariate analyses, not even cladistics. The data are analyzed to display patterns of similarity among the objects, and at least some of these patterns will reflect the history of the objects, but not necessarily in any explicit way. So, the fact that Ernst Mayr and Willi Hennig, for example, might have been familiar with the ideas of Charles Darwin, but not vice versa, is irrelevant to the analyses — all of the studied works are treated as equal.

The primary objective of the present paper is to demonstrate that this approach is equally applicable to humanities (e.g., Behrens and Yu 2003), including the historical sciences. We show that the simultaneous use of alternative procedures of exploratory data analysis may provide different insights into the same problem. In this way, we are able to reveal a pattern that was not disclosed by FL, and thus showing future directions towards an even more objective and meaningful evaluation of the history of thought in the biological sciences.

## 2 Methods

In the present study, we use exactly the same data as used in Fisler and Lecointre (2013, their Table 3): 41 works (OTUs^2^) described in terms of 91 variables, all of them nominal, with mostly 2, or rarely 3, states (possible values). Nominal variables represent the simplest type of data we can have: by using them the only judgment we can make about the OTUs is whether or not they possess the given variable. Ordering and differences between the possible states of the variable convey no meaning whatsoever. For example, in a given tree diagram drawn by some biologist the vertical axis may correspond to time (coded by 1) or not (coded by 0), as expressed by variable 38 of FL.

Our approach is dissimilarity-based, which means that the OTUs are compared in every possible pair by an appropriate mathematical function. The literature abounds in such measures, but in the present case our choice was limited: the data set contained many irrelevant or missing scores (there were 788=21% such entries in the matrix used by FL), which cannot be handled by most dissimilarity measures. We therefore used the Gower (1971) formula which can also handle variables of the nominal type. The formula takes the following form:

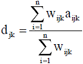

where n is the number of variables, a_ijk_ = 0 if OTUs j and k agree in variable i, and a_ijk_ = 1 otherwise. Weight w_ijk_ = 1 if OTUs j and k are comparable for variable i, and w_ijk_ = 0 if either or both OTUs have a missing or undefined score for that variable. The dissimilarity values have the range from zero to unity, 0 meaning complete identity and 1 referring to maximum dissimilarity.

All pairwise comparisons yielded a 41 × 41 dissimilarity matrix of OTUs, which was the starting point for all subsequent analyses, to produce a phenetic dendrogram (UPGMA clustering), a minimum spanning tree and a rooted additive tree (neighbor joining), a plexus graph and a phylogenetic network (neighbor net), and two ordinations (multidimensional scaling). Some comments on each of these methods will be given in the Results section, where the reader is referred to the cited literature on multivariate analysis and systematics for more details. The diagrams thus obtained are compared with each other and with the FL cladogram (called *Tree 1* in this paper; Figure 1) in order to determine whether: i) the cladogram nodes they recognized as meaningful indicators of groups (or schools, alternative approaches) are corroborated, and ii) any new information is also recovered from the data.

**Fig. 1.**
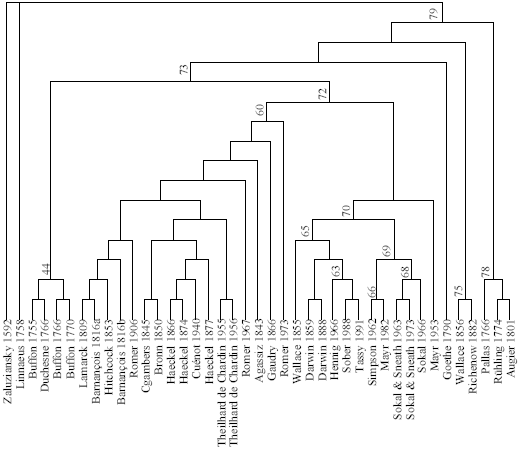
Consensus cladogram (Tree 1) of 279 equally parsimonious trees (378 steps) for 41 writings on trees and classifications in systematics. The tree is not drawn to scale, and only the sister group relations matter. Labels indicate “clades” recognized by the original authors and referenced here as well. Modified from Fisler and Lecointre (2013)

It is also important to note that these analyses place OTUs as sisters to each other, rather than placing some of them as ancestors and descendants, as would be true in an explicitly time-constrained analysis. It is impossible to determine from the data whether one OTU is the ancestor of another, and so they are placed in sister-group relationships.

Zaluziansky and Linnaeus are handled in the same manner as every other historical figure in all but one of the analyses. The exception is neighbor joining, in which Zaluziansky took a special position as an “outgroup” (see details below). Calculations were made using the SYN-TAX 2000 package (Podani 2001), except for the plexus graph drawn by the UciNet software (Borgatti et al. 2002) and the phylogenetic network computed by SplitsTree 4 (Huson and Bryant 2006).

## 3 Results

### 3.1 Tree 2: dendrogram

The phenetic alternative to conventional cladograms is the dendrogram, which converts dissimilarities to ultrametric distances (Lapointe and Legendre 1995). We used the group average (or UPGMA^3^, Sneath and Sokal 1973) algorithm for clustering, because it is also well-known in phylogenetic systematics, as a standard distance-based tree generating routine (meaningful whenever the molecular clock is “on”; Swofford et al. 1996; Page and Holmes 1998) and has been the most extensively used clustering procedure in many areas of science outside biology. For example, Babitch and Lebrun (1989) used this method for classifying languages and dialects, while Prieto et al. (2014) compared archaeological findings, namely terra-cotta figurines, by UPGMA. A dendrogram may be interpreted as a series of partitions (i.e., classifications into disjoint sets) in which small subsets (groups or clusters) are successively nested within large ones. The dendrogram may be “cut” at a given level to obtain a partition set.

Here, we recognized a partitioning into three major clusters (Figure 2, see also the Electronic Appendix), none of them in complete agreement with the “clades” in FL. In cluster A (“*Classifiers*”), which is the first one separated from the rest, we find tree users from the pre-evolutionary age of biology, plus some later authors who deliberately created a tree-based classification without evolutionary considerations (Wallace56 and Richenow, node 75 in FL). Goethe does not belong to this cluster, because he forms a singleton group, if we cut the tree around the dissimilarity level of 0.39. This reflects the ambiguity in his controversial views on “metamorphosis”, a fact still subject to intensive debate among historians of evolutionary biology (see e.g., Richards 2015; Spahn 2015). The special position of Goethe among the writers evaluated here is confirmed by the fact that in order to encounter the next singleton cluster (Haeckel66) one has to move down to a dissimilarity of 0.26. Note that on the FL cladogram, Goethe was also uniquely positioned.

**Fig. 2.**
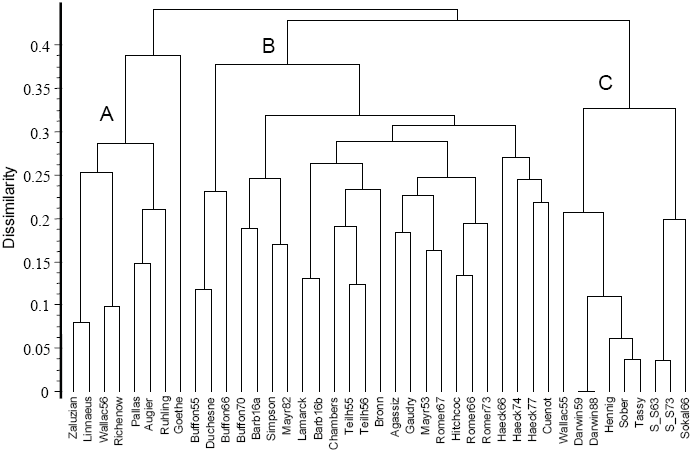
Group average (UPGMA) clustering (Tree 2) based on Gower dissimilarities of 41 writings on trees and classifications in systematics. Original labels are trimmed to 8 characters, and are still self-explanatory (but see Figure 1 or Figure 6 for full names, as used in FL). Letters identify three major groups, whereas Goethe remains as a singleton

Clusters B and C together (=FL node 73) include almost exclusively authors and works that recognized evolution, with different emphasis on its various aspects. The only exception is perhaps Agassiz, a believer in creation, whose presence in cluster B is due probably to the fact that he arranged fossils on the tree according to geological time. Cluster B unites the large group of metaphoricians (FL node 60) with Buffonians (FL node 44) and gradists (Mayr53, Mayr82 and Simpson – the latter two being FL node 66 as “grade theoreticians”). It is reasonable to call this large group collectively as “*Non-analytical evolutionists*” because subjective judgment had a primary role in their thinking about systematics. In their views, classifications enjoyed in most cases priority and evolution was considered only later to explain the classification.

Cluster C, on the other hand, comprises “*Modelers*”, who explicitly used trees to demonstrate evolutionary processes (Wallace and Darwin) or computed the tree to provide a starting basis for an *a posteriori* classification (cladists, FL node 63, and pheneticists, FL node 68). The relative closeness of cladists and pheneticists in Tree 2 may be surprising to some people, but they agree in many features, especially in their ambition to place biological classification on objective foundations, both theoretically and empirically. Also, the complex association of Haeckel, Darwin and Lamarck in the analyses is interesting, because both Haeckel and Lamarck saw evolution as an inherently progressive process, whereas Darwin did not.

All of the seven *Classifiers* are entirely homogeneous for five characters (emphasized by rectangles in the Electronic Appendix): 43 (state 0, the tree has classificatory aim), 44 (0, classification not made before the tree), 49 (0, the tree is not genealogical), 60 (0, time not considered) and 83 (0, no parsimony) – the latter two are also true for Goethe as well. There is no character state which would exclusively occur here. The relatively large group of *Non-analytical evolutionists* has only a single homogeneous variable, 83 (0, no parsimony). However, character states that predominate in this group, with no more than 3 exceptions or missing values, include: 1 (0, concrete ancestor at the root), 2 (0, no initial character states at the root), 3 (0, no inorganic forms included), 13 (0, no conceptual nodes), 35 (0, diversification axis carries no time), 47 (1, Nature is fundamentally ordered), 48 (0, tree is explicit), 51 (1, gradation in perfection), 72 (0, groups are not made according to genealogical links), 76 (1, groups are linked or nested), 84 (0, classification includes lack of shared properties), 86 (0, classification by global similarity) and 90 (0, homoplasies cannot be detected). Overall, cluster C, the *Modelers*, are the most homogeneous: they completely agree in 22 variables, and in a further 17 if Wallace55 is not considered (therefore the long list of variables is not given here). Not surprisingly, parsimony is the only character state that occurs exclusively in cluster C. See the Electronic Appendix for these character distributions.

### 3.2 Tree 3: minimum spanning tree

This tree connects OTUs directly such that the sum of weights assigned to the links (i.e. dissimilarities in this case) is the minimum (Rohlf 1973). It follows that terminal objects are always linked to their nearest neighbors. Successive removal of the longest links produces a hierarchical classification identical to the single-link clustering result. This type of tree was occasionally used in the initial period of numerical cladistics as a starting graph for phylogeny reconstruction. A noted example of using such trees outside biology is provided by Hage et al. (1996) in archaeology. In general, it serves as an alternative display of relationships to confirm or reject hypotheses of topological relationships. In our case, the overall arrangement of OTUs (Figure 3) reflects quite well the clusters of *Tree2*: the three groups are easily distinguishable along the main axis of the tree, which represents the longest path, that between Linnaeus and Sokal66. The largest dissimilarity separates group A from B (connecting Rühling with Barbançois16a), while the second longest edge separates Goethe from his nearest neighbor, Richenow, confirming the ambiguity of categorizing Goethe’s writings. Apparently, Mayr62 and Simpson, the theoretical gradists in FL, represent a somewhat transitional position between *Non-analytical evolutionists* (B) and *Modelers* (C). Buffon and Duchesne form their own subtree, the Buffonians (comparable with FL node 44). The metaphoricians (FL node 60) take the central position, from Barbançois16b to Agassiz. That is, in many details this tree agrees fairly well with *Tree 1* as well.

**Fig. 3.**
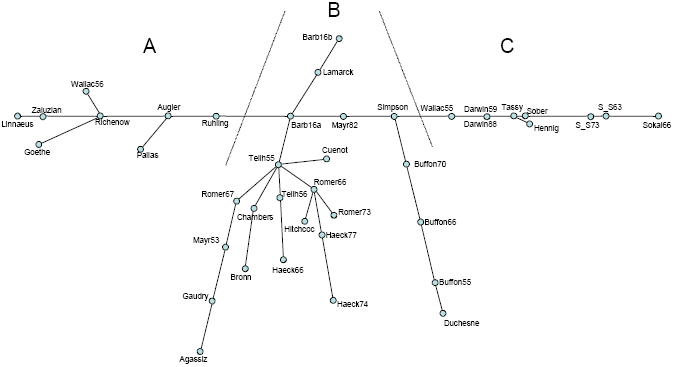
Minimum spanning tree (Tree 3) based on Gower dissimilarities of 41 writings on trees and classifications in systematics. Edge lengths are proportional to actual dissimilarities. Letters identify three major groups separated by dotted lines. Labels as in Figure 1

### 3.3 Tree 4: additive tree

The objective here is to generate a tree in which the between-object dissimilarities are as close as possible to the dissimilarities in the input matrix, and so clusters are not optimized directly. In this sense, this construct, most easily computed by the neighbor joining algorithm (Saitou and Nei 1987), is conceptually closest to cladograms, and it is often used in phylogenetics when the input matrix represents meaningful evolutionary distances. It has also been recommended as an adequate representation of manuscript traditions (Najock 1989). The algorithm produces an unrooted tree, which may be rooted by designating one OTU as the outgroup (here Zaluziansky) for comparability with rooted cladograms and dendrograms. As seen (Figure 4), Linnaeus is very close to Zaluziansky, justifying the decision of FL to select both of them as outgroups in parsimony analysis. Since it is not a clustering method, the large UPGMA groups are broken into parts that separate from the rest one by one as we proceed farther and farther from the root. The classifiers appear in two subtrees, with Goethe linked to Richenow and Wallace56 (see *Tree 3)*. It is remarkable that pheneticists (FL node 68) are separated from the strictly genealogical classifiers (FL node 65, from Wallace55 to Tassy), with some classifiers (FL node 78, e.g., Augier) and two gradists in between. This arrangement confirms the earlier findings that Mayr82 and Simpson are in a fairly equivocal position. The additive tree agrees with *Tree 1*, in that the Buffonians form a separate “clade” and that the metaphoricians (FL node 60) appear as an intact group.

**Fig. 4.**
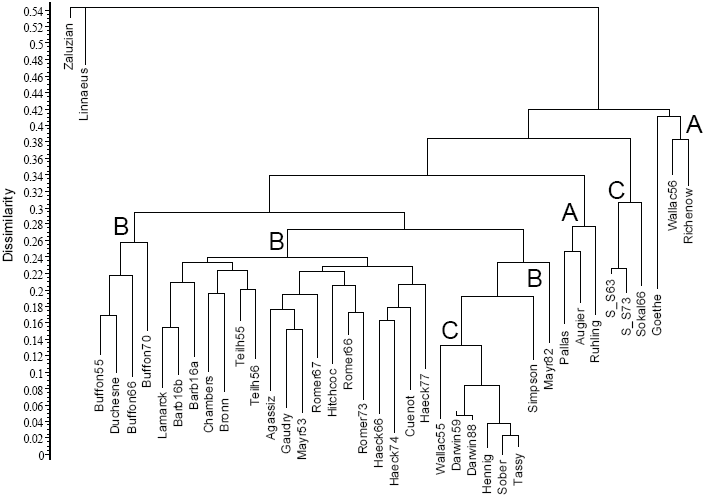
Additive tree representation (Tree 4) of Gower dissimilarities between 41 writings on trees and classifications in systematics. The outgroup is Zaluziansky. Letters refer to clusters identified in Figure 2, as broken into several subtrees here

### 3.4 Plexus graph

A conventional network graph differs from the minimum spanning tree in that there may be several different paths between two OTUs, i.e., there can be circular paths. Such graphs have been extensively used in the historical sciences (Gould 1993) and in citation analysis (Cronin and Atkins 2000). An example relevant to our study is provided by Krischel and Fangerau (2013), who compiled a social network for nineteenth century evolutionists, anthropologists and linguists, in which node size was determined by connectedness – Darwin’s node being the largest (their Figure 5). Such a graph is not appropriate here, however, because the relationship between writers is not of the yes-or-no type, but instead is measured on a continuous scale.

**Fig. 5.**
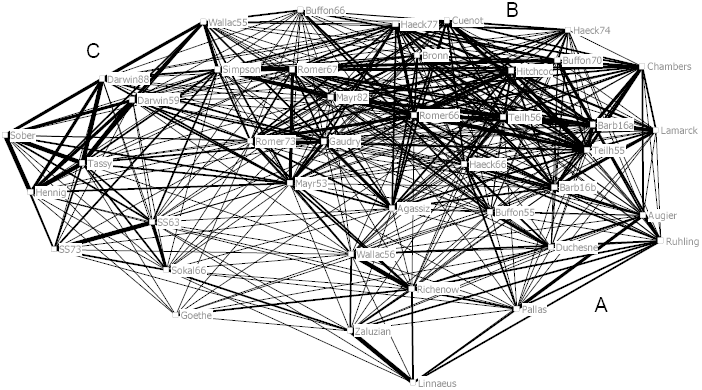
Plexus graph based on Gower dissimilarities of 41 writings on trees and classifications in systematics. See the text for an explanation of the line thickness categories. Letters refer to clusters identified in Figure 2

Therefore, we used a *plexus graph* in which the edges are drawn with different thickness (or color) depending on the dissimilarity between pairs of OTUs, a tool favored in the pioneering age of numerical ecology (McIntosh 1978). Here, we decided to use categories of dissimilarities, which are usually sufficient to reveal “coalitions” among the OTUs. These categories are, in decreasing order of line thickness: 0 ≤ *d* < 0.1; 0.1 ≤ *d* < 0.2; 0.2 ≤ *d* < 0.3, 0.3 ≤ *d* < 0.4. Pairs of writers with a dissimilarity of *d* ≥ 0.4 are not connected. The OTUs were arranged in a plane using the spring embedding algorithm. The plexus graph thus obtained confirms the existence of the three major groups recognized above (Figure 5). Most of the thickest edges connect members of *Modelers* (group C), which are associated to the *Non-analytical evolutionists* (group B) through weaker links (0.2 ≤ *d* < 0.3), with the exception of the connection between Wallace55 and Simpson. The cohesion within the *Non-analytical evolutionists* is weaker, whereas connectedness is fairly high, as it is within the *Classifiers* (group A). In the latter, Zaluziansky and Linnaeus, as well as Wallace56 and Richenow form close pairs. Duchesne and Augier represent the transition between Buffonians and the *Classifiers*. Note the central position of Mayr53, with links to all the three groups, and that of Goethe, who is apparently an outlier in the system.

### 3.5 Phylogenetic network

In addition to plexus graphs, there are many other types of networks used in biology. Those of particular interest here combine the hierarchical grouping properties of the clustering methods (see above) with the spatial representation of ordinations (see below) (Morrison 2014). These so-called “phylogenetic network” methods are increasing in popularity because they help test whether the data contain a strong tree-like signal, and will display a set of overlapping clusters if they do not. Note that the plexus network connects the OTUs via observed links, while phylogenetic networks connect them via inferred links and inferred nodes. The latter networks may be either more or less complex than the former. The use of such networks is by no means restricted to evolutionary biology (see Morrison 2014, for examples from other fields such as stemmatology, linguistics and archaeology). The main conceptual difference from trees is that trees produce nested groups whereas networks produce overlapping (i.e. non-exclusive) groups.

The neighbor net method, used here, starts from a dissimilarity matrix directly, producing a planar representation of the multivariate patterns. The resulting network (Figure 6) successfully displays 88.9% of the information in the original distance matrix. This is not a very tree-like network, indicating that the tree-based methods may be over-interpreting the groupings of the OTUs. Indeed, the phylogenetic network has more similarity to the ordination diagrams (see below) than to the trees (see above). The *Classifiers* and *Modelers* can be readily separated, but the *Non-analytical evolutionists* form a grade between them, as in *Tree 3* (Figure 3), with the Buffonians distinct from the rest. Goethe has a long terminal edge, as expected to indicate his equivocal position, but the gradists do not have an especially marginal position in the network. On the other hand, the three works by Haeckel are not closely associated in the network, which they are in the trees and also to some extent in the ordinations — this seems to reflect the complex patterns of missing data for these three works.

**Fig. 6.**
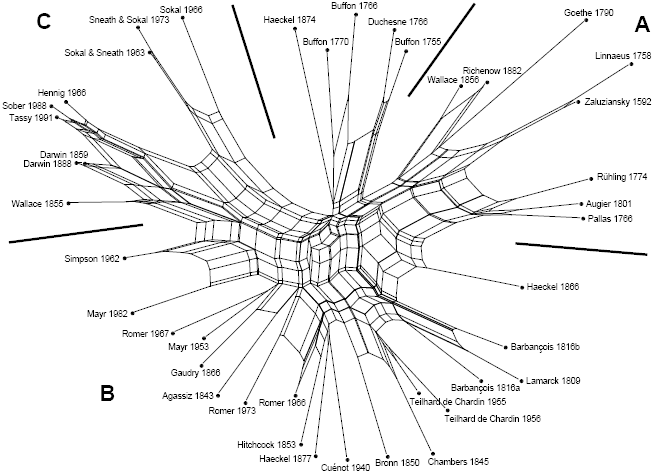
Neighbor network based on Gower dissimilarities of 41 writings on trees and classifications in systematics. Edge lengths are proportional to the original matrix distances. Letters identify three major groups separated by solid lines. The original labels used in FL are shown in full

### 3.6 Ordinations

As a supplementary tool for the line-graph representations, it is always worth trying some methods of ordination to reduce dimensionality in the original data space into a few axes (represented as a scatter plot), and then to evaluate whether clusters are distinguishable along these dimensions (Podani 2000). If the data set has a meaningful pattern because the original variables are correlated, then 2–3 ordination axes may be sufficient to display the inter-point relationships with a negligible loss of information. The “success” of the axes is expressed in terms of the percentage of eigenvalues of the starting matrix. Ordinations have been rarely used for phylogenetic purposes, but they are common in other fields of biology such as ecology, as well as in the archaeological sciences (Hodson et al. 1971). Since our raw data include too many missing values, only one group of ordination procedures is applicable here, namely multidimensional scaling, as these methods start from a dissimilarity matrix directly, in our case from the Gower dissimilarities.

We first used Principal coordinates analysis (PCoA), a metric procedure which arranges the OTUs in a new coordinate system such that the inter-point dissimilarities reproduce the original dissimilarities. Although no compact groups of OTUs are indicated (Figure 7), the arrangement of points is in complete harmony with the groups in *Tree2. Classifiers, Non-analytical evolutionists* and *Modelers* can be readily separated by straight lines in the first two dimensions. Goethe falls far from all other writers in the scatter plot, while the gradists Simpson and Mayr53 (but interestingly not Mayr82) take a marginal position in the group of *Non-analytical evolutionists*. Minor groups, such as pheneticists and cladists, are clear-cut in the diagram. The first eigenvalue explains 27.1% of the total variance, while the second one accounts for a further 21.6%, which at first glance suggests high explanatory power in these 2 dimensions. However, due to the often large and varying numbers of missing scores in the pairwise comparisons, there are many negative eigenvalues, with a total cumulative variance approximating 20% of the sum of positive eigenvalues.

**Fig. 7.**
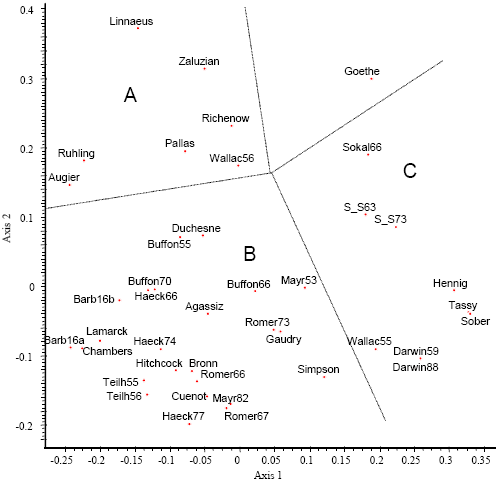
Principal coordinates ordination based on Gower dissimilarities of 41 writings on trees and classifications in systematics. Letters refer to clusters identified in Figure 2

The appearance of negative eigenvalues in the PCoA solution is indicative of the absence of true metric structure in the data, and the results may be doubtful in such cases. Thus, nonmetric multidimensional scaling is called for to confirm the picture obtained by PCoA. This arranges the OTUs in a pre-specified number of dimensions (usually two, representing a plane) such that the rank order of interpoint distances in the ordination is as close as possible to the rank order of the original dissimilarities. The analysis is iterative, by optimizing a random starting configuration; and the success of fit of the two rank orders is measured by the stress function, ranging from 0 to 1. We ran the program 20 times, and obtained the best result 3 times, with a stress of 0.207 – which is reasonable for 41 OTUs. The ordination (Figure 8) agrees with the PCoA result remarkably well suggesting that the lack of metric properties does not influence our conclusions regarding the groups. The major groups may be recognized as above, with Goethe isolated as always, and the gradists are again in a marginal position.

**Fig. 8.**
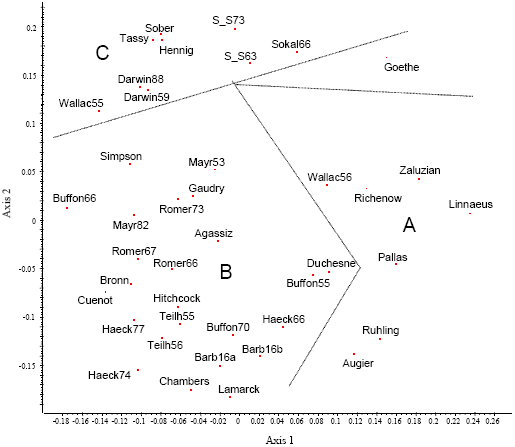
Nonmetric multidimensional scaling ordination based on Gower dissimilarities of 41 writings on trees and classifications in systematics. Letters refer to clusters identified in Figure 2; compare with the PCoA ordination in Figure 7

## 4 Discussion

This study used the same data as Fisler and Lecointre (2013; FL), although we do not agree completely with their selection of either scientists or characters. While most authors were represented only once, several others appeared twice or even three times in the FL study. This produced redundancy for those authors whose views did not vary much through time, especially for Darwin and Sokal & Sneath, and to some extent for Romer, Barbançois and Teilhard de Chardin as well. On the other hand, many important contributors to the history of systematics who also suggested or produced tree, tree-like or network summaries of their classifications were overlooked. To mention a few: Pax, Naudin, Herdman, Bessey, Hallier, Takhtajan, Whittaker, Cronquist, Doolittle and Cavalier-Smith – along with the entire school of pattern cladistics. The 91 selected characters are not optimal either. Due to missing scores, eight of them were not meaningful for more than 20 scientists, while three writings had undefined characters for more than 50 of the variables. Some variables were redundant, while none of them expressed the important distinction between a tree (as in Hennig) and a network (as in Buffon), for example. The dataset could thus be improved, although this would require a considerable amount of careful extra work.

However, to allow a direct comparison with the results of FL and to demonstrate the utility of other exploratory methods, we decided not to introduce changes. In one sense it is thus good news that our results confirmed several findings made by Fisler and Lecointre (2013), especially regarding the choice of outgroups, and the presence of minor “clades”. Not surprisingly, our *Tree 4*, the additive one (which may also be conceived as a distance based cladogram, i.e. a phylogram), agrees the best with *Tree1* (the FL cladistic tree) by being able to detect identical “clades”: initial tree users (node 78), tree makers (79), cladists (63), pheneticists (68), Buffonians (44), metaphoricians (60) and strictly genealogical classifiers (65). *Tree2* also shows three of these nodes, but not nodes 79, 44 or 60, while also reproducing the grade theoreticians (FL node 66). Of the nodes recognized and discussed by FL, the evolutionists (72) and connected graph users (70) are not reproduced by our analyses, mostly due to the “misclassification” or displacement of a few writings only. Also, the group of similarity classifiers (69 = 66+68), which appears so clear-cut in *Tree1*, is refuted by all of our diagrams.

The overall picture of the data structure differs in our analyses compared to FL, however. Most of our results suggest and others confirm – or do not refute at least – the observation that the scientific writings may be categorized into three separable, though not overly compact groups. There are some transitions between these groups, and also people who fit into more than one group. This picture is definitely more realistic than a single categorization since scientific ideas are never developed in isolation, all authors may influence the works of later authors, some concepts are inherited by new schools, others revised and still others completely reformulated. In other words, there is considerable fuzziness in the data which is best revealed by alternative approaches.

Regarding historical time of first appearance, the group of *Classifiers* includes authors who were not (yet) influenced by evolutionary theory in making their classifications or trees (such as Linnaeus and Augier) or who deliberately ignored evolutionary considerations, such as Wallace, who is otherwise considered together with Darwin as the developer of the theory of evolution through natural selection. The second group, *Non-analytical evolutionists* comprises authors who first recognized the existence of temporal change in organismal life, from Buffon through Lamarck to Romer. Even Agassiz is here, because he recognized that the fossil record changes through time, even though he was not an evolutionist. Gradists take a marginal position in this group, with weak affinities to the third group. In this third group, the *Modelers*, evolutionary change is explained by theoretical models, and its pathways are reconstructed or its results are evaluated by objective methodology. That is, Darwin and Wallace are not too far from Hennig conceptually, and, despite some philosophical differences, they are fairly close to the school of numerical taxonomy as well.

Goethe is certainly a unique thinker, an “outlier” – without having a close relationship to any of these groups. Notwithstanding the difficulties with the choice of data, we suggest that the three-group classification of scientists is a meaningful summary of tree-thinking in biological classification. Additional studies, with an expanded set of writings and more variables involved, may provide further insight into and a deeper understanding of that history.

The present study supports the general view that for the evaluation of complex data without obvious *a priori* structure, such as the dataset used here, the combination of various multivariate techniques may extract much more information than can any one analysis alone. An advantage of using alternative methods is that details supported by most procedures may be considered as “valid” structural properties of the data, such as the existence of many small clusters of writings in this study. Furthermore, in this way the limitations of one procedure may be compensated for by another. Clusters that appeared fairly distinct in the UPGMA dendrogram, for example, proved to be less clear cut in the networks and the ordinations. Although Fisler and Lecointre (2013) were skeptical about the usefulness of networks for demonstrating changes of biological thought, we found them to be as meaningful as any tree or ordination scatter plot.

We have thus shown that a purely cladistic approach to a classification problem, in which historical factors play little or no role, may be supplemented effectively by the joint application of various tree-and network-generating methods as well as ordinations, all of which are absolutely free from the assumptions of cladism.

Neither the cladistic method nor any of our alternative analyses are explicitly historical — historical patterns will be included in the outcome but they will not necessarily be separable from patterns resulting from any other source. In this paper, we have addressed whether the groups of people are robust by using different methods (i.e., the patterns are model independent), but we have not explicitly tested whether they have historical meaning. We have thus set up a series of hypotheses (the groups), and we have suggested possible historical interpretations of these groups, and so these hypotheses can now be examined in more detail and formally tested. The latter is beyond our brief, however.

Identifying the specifically historical pattern is, of course, important, but this goes beyond the capabilities of any multivariate analysis. A much more detailed assessment of the data would be required, which could now be based on the preliminary hypotheses presented here. This would include more than solely mathematical analyses, such as a detailed evaluation of the context of the individual writings studied here, perhaps with the inclusion of an expanded set of writings, and even then this may not be achievable with this type of intellectual inquiry.

## Acknowledgements

We thank Ferenc Jordán for preparing the diagram of the plexus graph. We are grateful to the anonymous referees for the constructive criticism of the manuscript.

## Footnotes

1 Here we use quotation marks, because clade is generally understood as a group of objects with common ancestry (monophyly) whereas in FLs study “clades” do not necessarily satisfy this requirement. These are groups formally optimized by a cladistic method.

2 In the terminology of numerical taxonomy, an OTU (= “operational taxonomic unit”) represents an individual study object, in our case a specified scientific writing due to a specified author. Each OTU appears as a single vertex (or node) in tree-like diagrams or as a point in ordination scatter plots.

3 Unweighted pair-group method using arithmetic averages.

## Electronic Appendix

The data from Table 3 of Fisler and Lecointre (2013) rearranged to follow the three-cluster classification obtained by UPGMA clustering (groups A to C), such that the sequence of writings is in the same order as in the dendrogram of Fig. 2. Blocks of scores are emphasized by rectangles in order to illustrate variables that have the highest explanatory power.

